# Local structural dynamics of Rad51 protomers revealed by cryo-electron microscopy of Rad51-ssDNA filaments

**DOI:** 10.1101/2024.05.06.592824

**Authors:** Jie Liu, Steven Gore, Wolf-Dietrich Heyer

**Author notes:** To whom correspondence should be addressed: Jie Liu, Department of Microbiology & Molecular Genetics, University of California, Davis, One Shields Ave., Davis, California 95616-8665, Wolf-Dietrich Heyer, Department of Microbiology & Molecular Genetics, University of California, Davis, One Shields Ave., Davis, California 95616-8665.

## Abstract

Homologous recombination (HR) is a high-fidelity repair mechanism for double-strand breaks. Rad51 is the key enzyme that forms filaments on single-stranded DNA (ssDNA) to catalyze homology search and DNA strand exchange in recombinational DNA repair. In this study, we employed single-particle cryo-electron microscopy (cryo-EM) to ascertain the density map of the budding yeast Rad51-ssDNA filament bound to ADP-AlF_3_, achieving a resolution of 2.35 Å without imposing helical symmetry. The model assigned 6 Rad51 protomers, 24 nt of DNA, and 6 bound ADP-AlF_3_. It shows 6-fold symmetry implying monomeric building blocks, unlike the structure of the Rad51-I345T mutant filament with three-fold symmetry implying dimeric building blocks, for which the structural comparisons provide a satisfying mechanistic explanation. This image analysis enables comprehensive comparisons of individual Rad51 protomers within the filament and reveals local conformational movements of amino acid side chains. Notably, Arg293 in Loop1 adopts multiple conformations to facilitate Leu296 and Val331 in separating and twisting the DNA triplets. We also analyzed the predicted structures of yeast Rad51-K342E and two tumor-derived human RAD51 variants, RAD51-Q268P and RAD51-Q272L, using the Rad51-ssDNA structure from this study as a reference.

## INTRODUCTION

Double-strand breaks (DSBs), gaps, and interstrand crosslinks (ICLs) in DNA disrupt normal cellular functions and, if left unrepaired, can lead to cell death. HR is a highly conserved pathway that repairs such damages by utilizing an intact DNA template from the sister chromatid or homologous chromosome when available (1,2). DNA resection, homology search, DNA strand exchange, and new DNA synthesis encompass the central steps of HR, while subsequent steps determine the amount and configuration of new genetic information incorporated into the repaired DNA. Dozens of proteins are involved in HR process and regulation, but the key catalytic unit is the filament formed by the essential enzyme Rad51 on ssDNA which performs the signature reactions of homology search and DNA strand exchange (3,4).

Nucleotide cofactors directly impact the DNA-binding affinities of Rad51 through the highly conserved ATPase core domain of the Rad51 family of proteins (5). The ATP-bound form of Rad51 displays high affinity towards ssDNA and forms an “active” nucleoprotein filament by extending B-form-like ssDNA about 1.5 fold to reach a helical pitch of ∼ 95-100 Å (6,7). The ADP-bound form of Rad51 forms an “inactive” nucleoprotein filament in a compressed state with a helical pitch of ∼ 75-85 Å (7,8). The presence of ATP enhances the assembly of Rad51 filaments on DNA, resulting in enhanced stability and resistance to nuclease digestion (9,10). Even though ATP hydrolysis by Rad51 is dispensable for its DNA strand exchange activity, it is important to maintain a stable and readily available protein pool of Rad51. Turnover of ATP and, consequently, Rad51 filaments prevents the formation of overly stable Rad51 filaments and depletion of free Rad51 *in vivo* (11–14). Likewise, DNA binding has an influence over how Rad51 interacts with ATP. Wild type Rad51 alone shows minimal ATPase activity, similar to the Rad51-K191A or -K191R mutants in the Walker A box of its ATP binding site. The presence of DNA stimulates the ATP hydrolysis activity of the wild type Rad51 protein significantly, but it has no effect on the activity of the Rad51-K191A and -K191R mutants (9,15). Thus, the interactions of Rad51 with ATP and DNA are interconnected and impactful (5,11).

Structures of bacteria, archaea, yeast, and human homologs of Rad51 filaments on DNA show highly conserved core domains responsible for ATP, DNA, and protomer-protomer interactions (16–19). The structures of Rad51-I345T and Rad51-H352Y, two yeast Rad51 mutant proteins that exhibit enhanced DNA-binding activities, have been solved by X-ray crystallography (18,20). Rad51-I345T partially bypasses the requirement of Rad51 paralogs, Rad55 and Rad57, through its increased affinity for DNA, even in the presence of RPA (21). His352 is a key residue at the Rad51 protomer-protomer interface. The Rad51-H352Y mutant forms stable filaments on ssDNA with enhanced resistance to salt disruption but lacks ATP hydrolysis and DNA strand exchange activities (20,22). Surprisingly, both Rad51 mutant structures lack DNA and nucleotide cofactors, even though they bind DNA more tightly than the wild type protein. Additionally, the Rad51-I345T filament exhibits a three-fold symmetry with a dimeric building block, while the Rad51-H352Y filament presents a six-fold symmetry and uniform protomer-protomer interfaces with a monomeric assembly unit. A high-resolution structure of wild type yeast Rad51 filaments with discernible DNA and nucleotide cofactors is needed to resolve conflicts of the Rad51-Rad51 interface and advance our understanding of the interplay between ATP and DNA and their impact on Rad51 conformation.

RAD51-Q268P and RAD51-Q272L are deleterious RAD51 mutants detected in human lung adenocarcinoma and kidney carcinoma tumors, respectively (23). A thorough biochemical analysis has revealed that both mutants exhibit decreased affinity for ssDNA and dsDNA binding, with Q272L showing more severe defects. Interestingly, both mutants have lost significant activities in basal level and ssDNA-stimulated ATP hydrolysis as compared to the wild type protein. Q268P was found to be unable to catalyze a DNA strand exchange assay with phage DNA substrates, whereas Q272L retained some activity (23). Q268P and Q272L are located in the DNA-binding Loop2 and away from the ATP-binding pocket, and more structural analysis is required to undertand the mechanisms underlying the observed biochemical defects.

Cryo-EM has emerged as a promising technique for achieving atomic resolution in the study of macromolecules. Compared to traditional X-ray crystallography methods, cryo-EM offers the advantage of preserving macromolecules in a more native environment. In this study, we utilized cryo-EM single-particle image analysis to obtain a high-resolution structure of a budding yeast Rad51 filament bound to ssDNA and the non-hydrolysable transition-state ATP analog ADP-AlF_3_ without imposing helical symmetry. Our analysis of the Rad51 helical filament provides insights into the conformational dynamics of amino acid side chains through the comparison of individual Rad51 protomers within the filament. Notably, we observed movements of Ser192 and Arg293 associated with ATP hydrolysis and DNA binding, as well as amino acids involved in protein binding. Furthermore, we integrated molecular modeling with AlphaFold predictions to elucidate the structural basis for the altered activities of Rad51 mutants, including yeast Rad51-I345T and Rad51-K342E, as well as human RAD51-Q268P and RAD51-Q272L.

## MATERIALS AND METHODS

### DNA and protein purification

The 5’-biotinylated 100mer used in this study was described previously with the following sequence, 5’-biotinylated AACGACGTTTGGTCAGTTCCATCAACATCATAGCCAGATGCCCAGAGATTAGAGCGCATGACAAGTAAAGGACGGTTGTCAGCGTCATAAGAGGTTTTAC-3’ (24). Native untagged budding yeast Rad51 protein was purified as described (25,26).

### DNA-binding assay

15 μL reactions containing 0, 250, 500, 1,000, and 2,000 nM Rad51 were incubated with 3 μM nt ϕX174 ssDNA (5,386 nt; NEB) in buffer containing 2 mM nucleotide (where indicated), 35 mM Tris-acetate pH 7.5, 7 mM magnesium acetate, 50 mM NaCl, 1 mg/mL BSA, and 1 mM DTT for 30 min at 30 °C. Reaction products were separated on 0.7% agarose gel in TA buffer (40 mM Tris-acetate pH 8) at 4 °C and visualized by Sybr Gold staining (Thermo Fisher Scientific).

### Cryo-EM sample preparation and data acquisition

15 μM Rad51 was incubated with 45 μM nt of the 5’-biotinylated 100mer in in buffer containing 35 mM Tris-acetate pH 7.5, 7 mM magnesium acetate, 50 mM NaCl, 5% glycerol, 1 mM ADP-AlF_3_, and 1 mM DTT for 15 min at 30 °C to form nucleoprotein filaments before freezing. 5 μL of Rad51-ssDNA complexes was deposited on thin carbon-coated Quantifoil grids and incubated for 1 min before vitrification using a Vitrobot Mark IV (Thermo Fisher Scientific). Cryo-grids was initially screened using a Thermo Fisher Glacios equipped with Gatan K3 direct electron detector. A large-scale cryo-dataset was collected using a Titan Krios microscope (Thermo Fisher Scientific) equipped with a K3 detector and a BioQuantum energy filter at the Stanford-SLAC Cryo-EM Center (S^2^C^2^) (27). 25,659 movies were collected using SerialEM at a nominal magnification of 130,000, corresponding to a pixel size of 0.68 Å. The movies were collected with an exposure mode of 40 frames in 1.85 seconds to reach a total dose of 60 e^-^/ Å^2^ and a defocus range of -0.5 to -2.5 μm.

### Cryo-EM image processing and 3D map reconstruction

The image processing procedure using CryoSPARC is summarized in **Supplementary Figure 1** (28). The original movies were imported into a CryoSPARC database for motion correction, and dose-weighted/non-dose-weighted micrographs were generated before the contrast transfer function (CTF) estimation. Low-quality micrographs were eliminated after manual inspection and particles were picked automatically with template picker from selected micrographs. Iterative rounds of 2D classification were performed to remove junk classes and associated poor-quality filaments. A final particle set containing 2,941,384 filaments were used to create three *ab initio* models for following 3D homogeneous refinement. After multiple runs of 3D refinement and CTF refinement, 1,901,160 out of 2,941,384 particles were included to reconstruct the final density map of Rad51-ssDNA filaments at a resolution of 2.35 Å based on the Fourier shell correlation (0.143 Å) criterion. A second species of Rad51 filaments were reconstructed to a 3D map at 2.52 Å, using 754,384 particles.

### Model building and refinement

The previously published yeast Rad51-I345T crystal structure was used to generate an initial atomic model that was docked into the EM map by using PHENIX (29). The ADP-AlF_3_, magnesium ion and ssDNA were manually fit into the density map as ligands in COOT (30,31). Based on the sequence-nonspecific DNA-binding behavior of Rad51, we used a poly-dT sequence to build the ssDNA in the atomic model. Multiple building and refinement were performed in PHENIX and COOT to improve model accuracy with no imposed symmetry. Final atomic models were visualized and analyzed in CHIMERA (32).

### Structural comparison and modeling

To create the model of the yeast Rad51-dsDNA filament structure, we superimposed the yeast Rad51-ssDNA model from this study with the human RAD51-dsDNA cryo-EM structure (7eje) in CHIMERA. Based on the almost perfect superimposition of the ssDNA in the yeast Rad51-ssDNA structure and one of the strands from the dsDNA in the human RAD51-dsDNA structure, we replaced the ssDNA with the dsDNA to create the yeast Rad51-dsDNA filament model.

The structural comparisons of the yeast wild type Rad51 structure (PDB:9B2D) with the Rad51-I345T (PDB: 1SZP) and Rad51-H352Y (PDB: 3LDA) mutant proteins were performed in CHIMERA. The structures of the yeast and human Rad51 mutants were predicted and generated by AlphaFold 2, which was further compared to the wild type Rad51 structure in CHIMERA.

### Sequence alignment

Alignments of Walker A and B nucleotide binding motifs, and the regions around Arg293 and Val331 were generated using Clustal Omega. Abbreviations: *Ec*: *Escherichia coli*, *Sc*: *Saccharomyces cerevisiae*, *Sp*: *Schizosaccharomyces pombe*, *At*: *Arabidopsis thaliana*, *Ce*: *Caenorhabditis elegans*, *Dm*: *Drosophila melanogaster*, *Dr*: *Danio rerio*, *Mm*: *Mus musculus*, *Hs*: *Homo sapiens*.

## RESULTS

### Cryo-EM structure of Rad51-ssDNA filament with ADP-AlF_3_

We purified yeast Rad51 protein and screened Rad51-ssDNA filament formation using different nucleotides (**Fig. 1A, B)**. To achieve optimal protein-DNA complex formation, we incubated Rad51 with a 100-mer ssDNA and ADP-AlF_3_, which enabled us to obtain high-quality micrographs for cryo-EM studies. Instead of conventional helical filament analysis, we employed single-particle image analysis with no imposed symmetry to obtain cryo-EM structures of the Rad51-ssDNA filaments in the presence of ADP-AlF_3_ at a resolutions of 2.35 Å and 2.52 Å (**Fig. 1C, Suppl. Fig. 1**). The parameters of data collection, refinement, and model validation were listed in **Supplementary Table 1**. Our study yielded two almost identical right-handed helical structures with a 100 Å pitch and ∼ 6.3 Rad51 protomers per turn (**Suppl. Fig. 2**). The reconstructed maps were subjected to an automatic symmetry search, which revealed the presence of 6.32 Rad51 protomers per turn in the higher-resolution map and 6.31 Rad51 protomers per turn in the lower-resolution map (**Suppl. Fig. 2**). The parameters of data collection, refinement, and model validation are listed in **Supplementary Table 1**. The spatial forms and helical parameters of our reconstructed maps are similar to those of previously published human RAD51-DNA cryo-EM structures (33,34).

**Figure 1.**
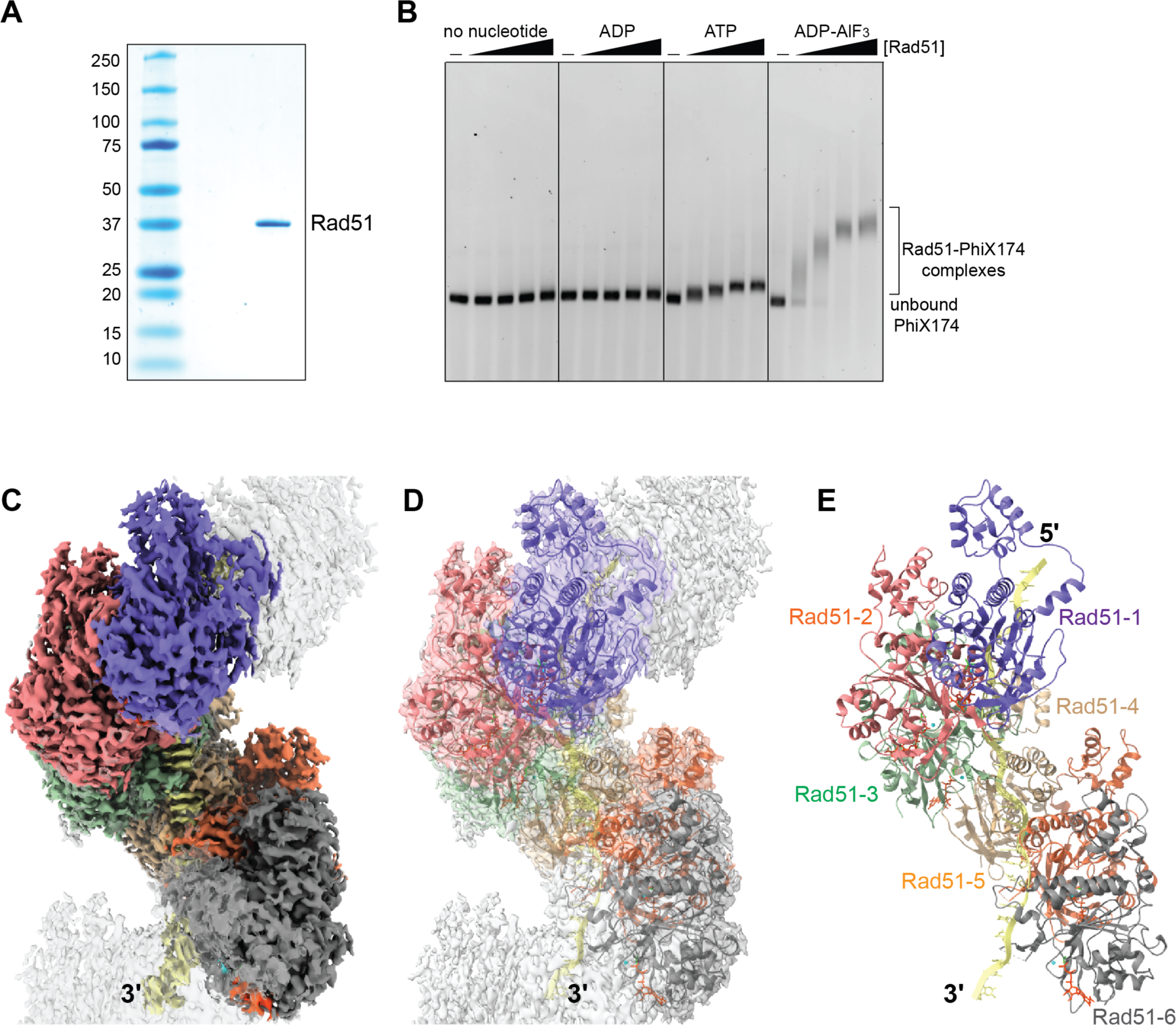
Cryo Structure of yeast Rad51-ssDNA filament with ADP-AlF_3_. (**A**) 1 µg of purified yeast Rad51 visualized on a 4-20% gradient SDS-PAGE stained with Denville Blue. Protein standards were labeled on the left. (**B**) EMSA of Rad51 binding on <X174 ssDNA with no nucleotide, 2 mM of ADP, ATP, or ADP-AlF_3_. (**C**) Density map of Rad51-ssDNA filament with six color-coded Rad51 protomers and ssDNA in yellow. (**D**) Overlay of density map in **C** with the molecular model in **E**. Rainbow-colored Rad51 protomers are individually labeled in **E**.

We built an atomic model of the Rad51-ssDNA filament using the 2.35 Å map, containing six Rad51 protomers that are bound to the nucleotide cofactor ADP-AlF_3_ and 24 nucleotides of ssDNA (**Fig. 1D, E**). The ADP-AlF_3_ and magnesium ions are positioned at the interface between two Rad51 protomers, surrounded by the Walker A and Walker B motifs from one Rad51 protomer and one alpha-helix (α15) and one loop from the neighboring Rad51 (**Fig. 2A, B**). The fits of the modeled ADP-AlF_3_, magnesium ion, and ssDNA nucleotides with the well-resolved density map are illustrated in **Figures 2C & G**. We were unable to observe the densities of the first 78 amino acids at the N-terminus and several amino acids in the flexible DNA-binding Loop2, indicating that these regions are highly dynamic. These observations are consistent with the yeast Rad51 crystal and human RAD51 cryo-EM structures (18,20,33,34).

**Figure 2.**
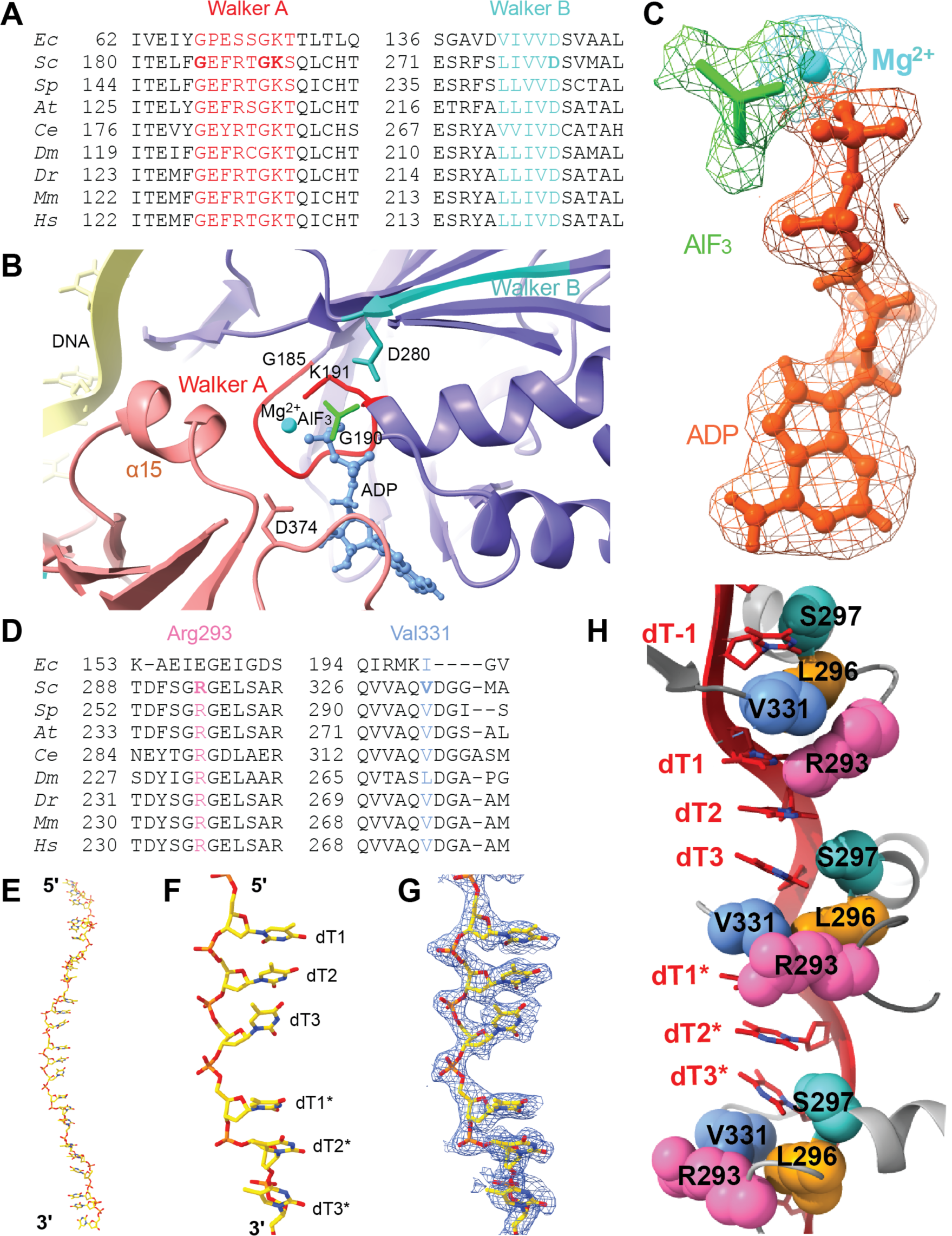
ADP-AlF_3_ and ssDNA binding sites. (**A**) Sequence alignment of Walker A and B nucleotide-binding motifs. **(B)** Positions of Walker A and Walker B motifs around ADP-AlF_3_ and Mg^2+^. (**C**) Quality of ADP-AlF_3_ fit to density map. (**D**) Sequence alignment of Arg293 and Val331 regions. (**E**) Modeled ssDNA. (**F**) Two ssDNA triplets showing the twist of the bases. (**G**) Quality of ssDNA fit to density map. (**H**) Role of Arg293, Leu296, Ser297, and Val331 in twisting and unstacking of the DNA bases.

The binding and stretching of ssDNA by Rad51, which occurs in a nucleotide triplet pattern (**Fig. 2E-H**), is similar to that of *E. coli* RecA and human RAD51 (33–35). Our analysis identified key amino acid residues within the DNA-binding Loop1, specifically Arg293, Leu296, and Ser297, as well as Val331 in the DNA-binding Loop2 of the neighboring Rad51, which play a crucial role in partitioning nucleotide triplets (**Fig. 2H**). These residues are highly conserved in eukaryotic Rad51 proteins (**Fig. 2D**). The positively charged channel, positioned along the interior of the Rad51 filament, binds to the negatively charged phosphodiester backbone. Additionally, Leu296 and Val331 from two adjacent Rad51 protomers were inserted between the nucleotide bases from neighboring triplets. These two bases are also pushed by Arg293 and Ser297, such as dT3 and dT1* in **Figure 2H**. The combined efforts of these four amino acids result in a separation of the triplets by breaking the base stacking, as illustrated in **Figure 2H** and **Supplemental Figure 3A, B**.

The yeast Rad51-I345T mutant filament exhibits a three-fold helical symmetry, as published previously, and comprises two types of protein-protein interfaces (18). These interfaces stem from two distinct Rad51-I345T protomers, which display slight movement and twist at the N-terminal domains using the C-terminal ATPase core domain as a reference (**Fig. 3A**, (18)). However, when all six Rad51 protomers from this study align for comparison, we do not observe different types of protein-protein interfaces (**Fig. 3B**). This finding is similar to the uniform protomer-protomer interface observed in the crystal structure of Rad51-H352Y (20). From these observations, we conclude that wild type Rad51 forms DNA filaments with single protomer building blocks and that the dimeric building block structure of the Rad51-I345T ssDNA filament is specific to this mutant (see also below).

**Figure 3.**
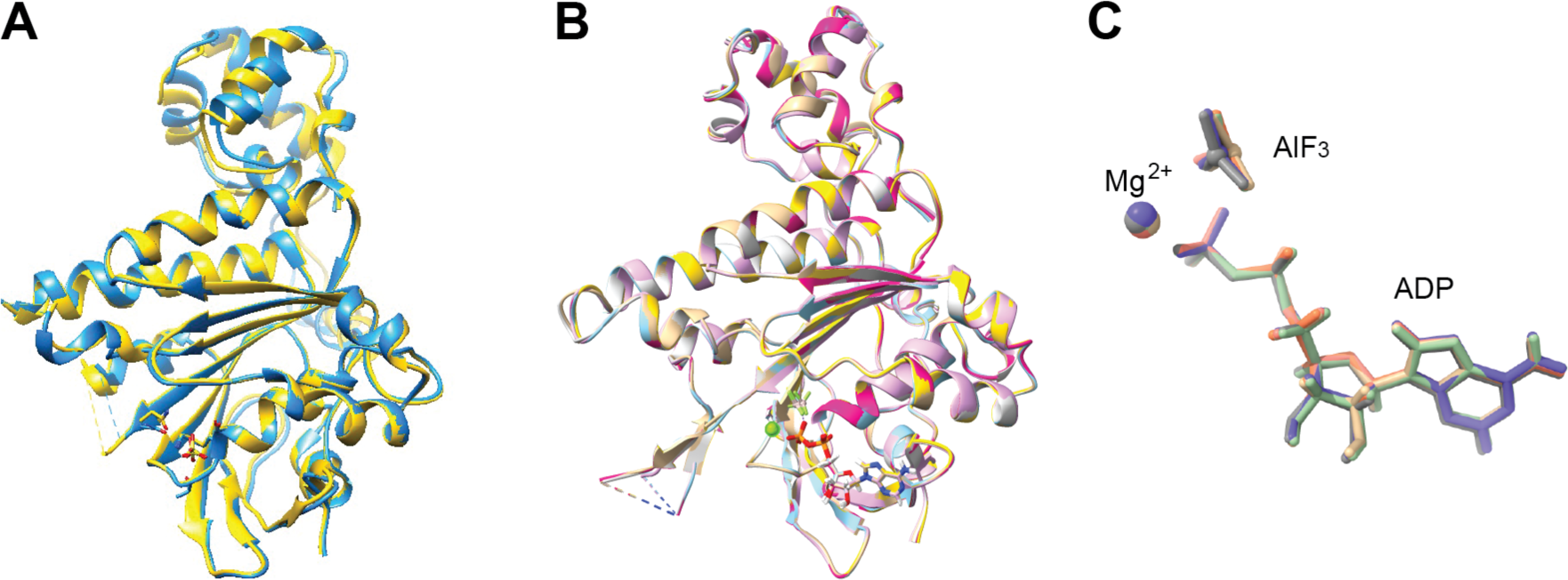
Identical binding interface and individual Rad51 protomers. (**A**) Overlay of Chain A and Chain D protomers of Rad51 from yeast Rad51 crystal structure (1SZP). (**B**) Overlay of six individual Rad51 protomers from Fig. 1E. (**C**) Enlarged view of bound ADP-AlF_3_ from **B**.

### Local dynamics of amino acids of Rad51 involved in ATP-binding

We noticed that the AlF_3_ were trapped at different positions, despite the remarkable similarity between ADP and the protein backbone in the superimposition of six Rad51 protomers (**Fig. 3C**). We entertained the idea of whether reaction intermediates could be captured by aligning all the six Rad51 protomers, with a focus on amino acid side chains. Indeed, we discovered the side chain of Ser192 could interact and pull AlF_3_ away from ADP (**Fig. 4A, B**). The varied positions of Ser192 and AlF_3_ suggest how the movement of Ser192 might work with the γ-phosphate to facilitate Lys191 in catalyzing ATP hydrolysis. Interestingly, Lys191 does not show any movement, but that may be due to the presence of the non-hydrolysable transition state analogue, ADP-AlF_3_, used in the filament assembly.

**Figure 4.**
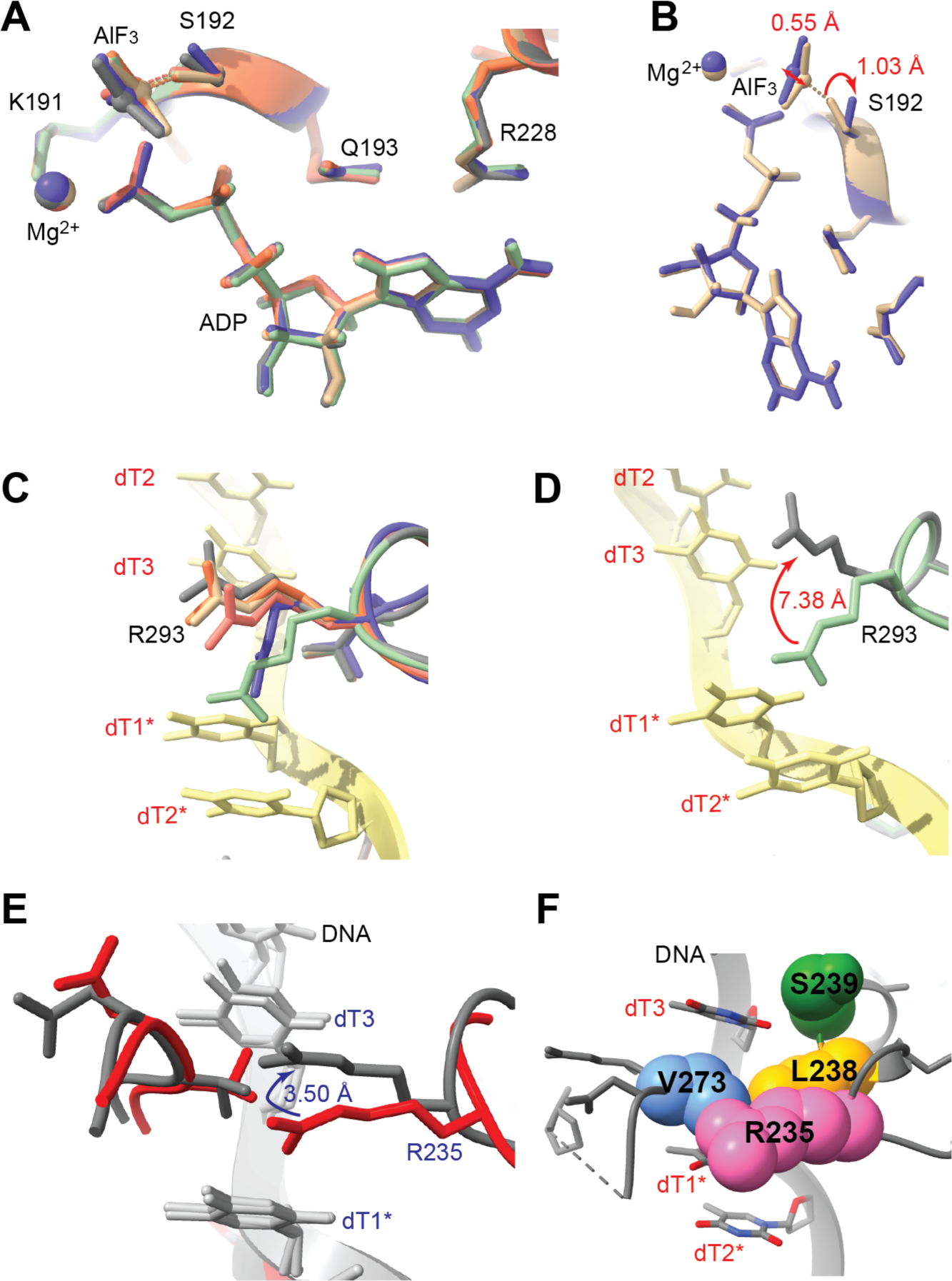
Movement of key residues during ATP hydrolysis and ssDNA binding. (**A**) Ser192 is pulling AlF_3_ away from ADP to facilitate ATP hydrolysis. (**B**) The maximal movement of Ser192 in two different Rad51 protomers is shown in purple and tan. (**C**) Multiple conformations of Arg293 in different Rad51 protomers and maximal distance shown in **D**. DNA are shown in yellow with bases labeled. Amino acids from each Rad51 protomer are labeled with different colors. (**E**) Positions of Arg235 in published cryo-EM structures of human RAD51-ssDNA filaments, shown in red (5h1b) and gray (7ejc). SsDNA is shown in light gray. (**F**) Role of Arg235, Leu238, Ser239, and Val273 in separating the DNA triplets.

The crystal structure of Rad51-H352Y was obtained with six-fold helical symmetry, but multiple conformations of amino acid side chains were still present. Specifically, it has been observed that the Tyr352 residue adopts two distinct conformations in a 1:1 ratio, as indicated by mixed density (20). We also examined published cryo-EM structures of human RAD51 structures with ssDNA or dsDNA, derived from imposing helical symmetry, based on the high structure and function conservation of yeast and human Rad51 proteins. Human Thr134 (yeast Ser192) also adapts different conformations, pulling Mg^2+^ and the γ-phosphate (**Suppl. Fig. 3C, D**) in a similar way to yeast Ser192 (**Fig. 4A, B**). The orientation of Lys133 remained unchanged as the equivalent yeast Lys191 (**Suppl. Fig. 3C,D**). This consistency confirms the local dynamics of residues in yeast and human Rad51 involved in ATP hydrolysis.

### Local dynamics of amino acids of Rad51 involved in DNA-binding

Next, our investigation delved into the local movements of the amino acids involved in DNA binding. As illustrated in **Figure 4C**, Arg293 from different yeast Rad51 protomers exhibits multiple conformations, distributed within the top and bottom nucleotide bases from two adjacent triplets (dT3 and dT1* in **Fig. 2H**). The different positions of Arg293 show a maximal distance of 7.38 Å and suggest a pushing action towards the nucleotide base dT1* (**Fig. 4D** & **2H**). Again, we compared four known human RAD51 filament cryo-EM structures. Arg235 from two different human RAD51-ssDNA structures (7ejc and 5h1b) also displayed two conformations within two nucleotide bases with a 3.5 Å distance (**Figure 4E**). It is worth noting that in the human RAD51-dsDNA structures, the side chains of Arg235 and Asp274 exhibit changes in orientation towards the complementary strand (**Suppl. Fig. 3E,F**). These consistent movements of yeast Arg293 and corresponding human Arg235 may play crucial roles in coordinating the insertion of Leu296 and Val331 (human Leu238 and Val273) to separate neighboring DNA triplets during the presynaptic stage while stabilizing complementary strands in the postsynaptic stage (**Fig. 2H** & **4F**).

### Local dynamics of amino acids of Rad51 involved in protein-interaction

We continued to investigate whether the amino acids involved in self-association or hetero-protein association also exhibit local movements. First, we analyzed the conformations of the amino acids in the α-helix (α6) located at the Rad51-Rad51 interaction interface (**Fig. 5A**). Of the residues examined, Asp149, Arg153, Glu156, and Ile158 exhibit multiple orientations, all positioned away from the filament axis and exposed to the solvent (**Fig. 5B**). Conversely, Phe144, Thr146, Ala147, and His151 interact with the neighboring Rad51 protomer and display only one orientation with minimal movements (**Fig. 5C**).

**Figure 5.**
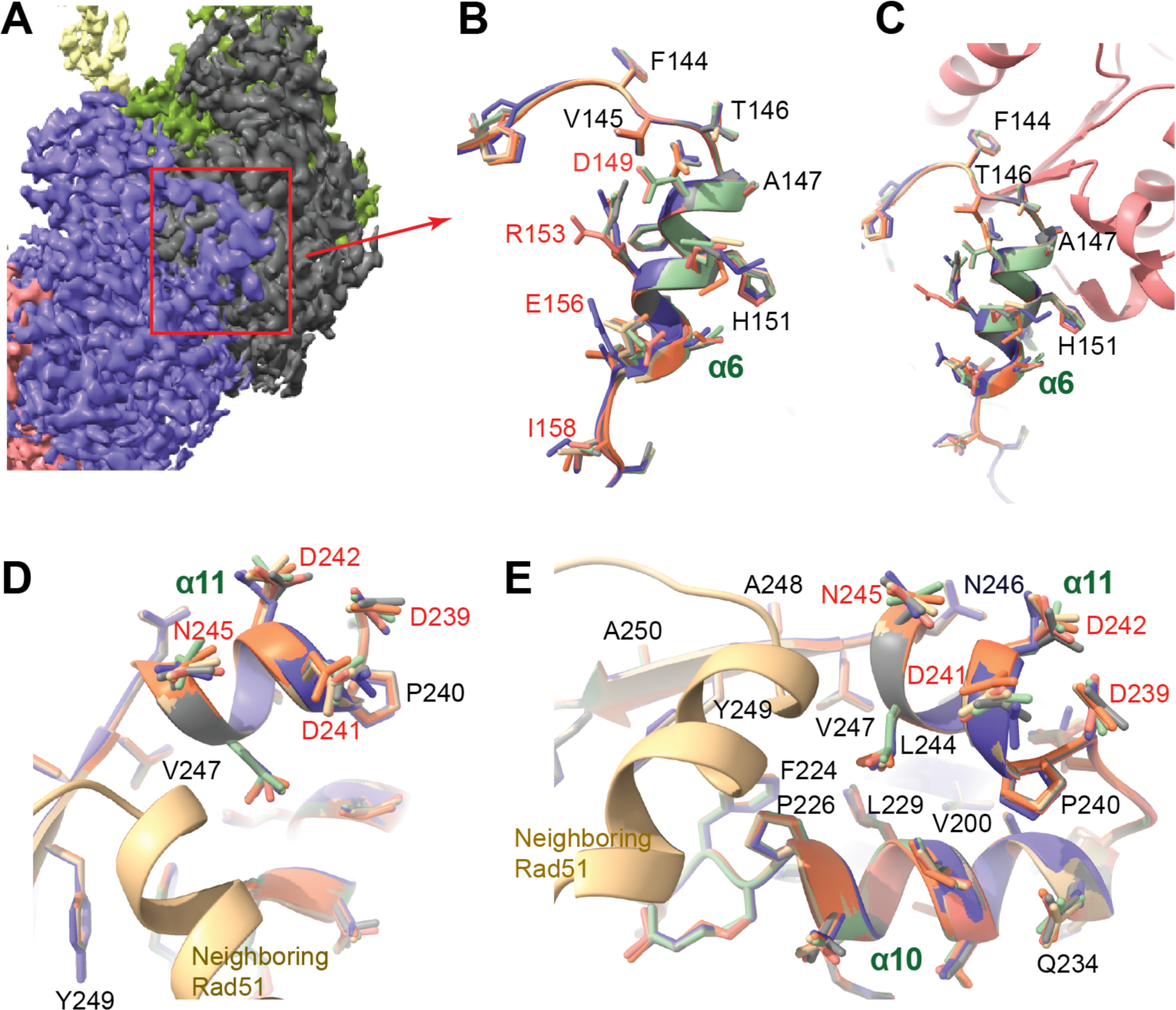
Solvent-exposed residues on alpha-helices at the protein-interaction interfaces are flexible, showing multiple conformations. (**A-C**) Conformations of flexible and rigid amino acids at the Rad51-Rad51 interface. The marked α-helix (α6) and adjacent loop in **A** were enlarged in **B** and **C**. (**D**) Multiple conformations of Asp239, Asp241, and Asp242 in the acidic patch (α11) involved in hetero-protein association. (**E**) Amino acids in the neighboring α-helix (α10) and β-sheet around the acidic patch show minimal movement. Rigid residues are labeled in black, while flexible ones are in red.

Next, we inspected the evolutionarily conserved acidic patch discovered in fission yeast Rad51, which is involved in hetero-protein associations with multiple partners (36). In budding yeast Rad51, this patch comprises three aspartates (Asp239, Asp241, Asp242) which are highly dynamic and are oriented towards the solvent (**Fig. 5D**), thereby accommodating a sensing configuration to search and interact with other recombination protein factors such as Rad52, Rad55-Rad57, and Rad54 (36). Our observations also reveal that the amino acids present on the neighboring β-sheets, such as Val247, Ala248, Tyr249, interact with the adjacent Rad51 without exhibiting any alternative conformations, irrespective of their facing direction (**Fig. 5E**). Similarly, the amino acids on the neighboring α-helix (α10), from Pro226 to Gln234, also interact with adjacent Rad51 and show minimal movement.

### A 3D model for Rad51 mutant analysis

To evaluate the Rad51 mutants with dsDNA-binding defects, we modeled a Rad51-dsDNA filament structure. First, the yeast Rad51 model was compared with the human RAD51-dsDNA cryo-EM structure (7eje), and the two structures superimposed well with almost identical protein-DNA contacts (**Fig. 6A**). Furthermore, the positions of ssDNA backbone and bases in yeast Rad51-ssDNA structure aligned almost perfectly with one strand from the human RAD51-dsDNA (**Fig. 6A, B**), enabling us to model a yeast Rad51-dsDNA filament (**Suppl. Fig. 4A, B**). This model holds significant implications for the analysis of Rad51 mutants with dsDNA-binding defects.

**Figure 6.**
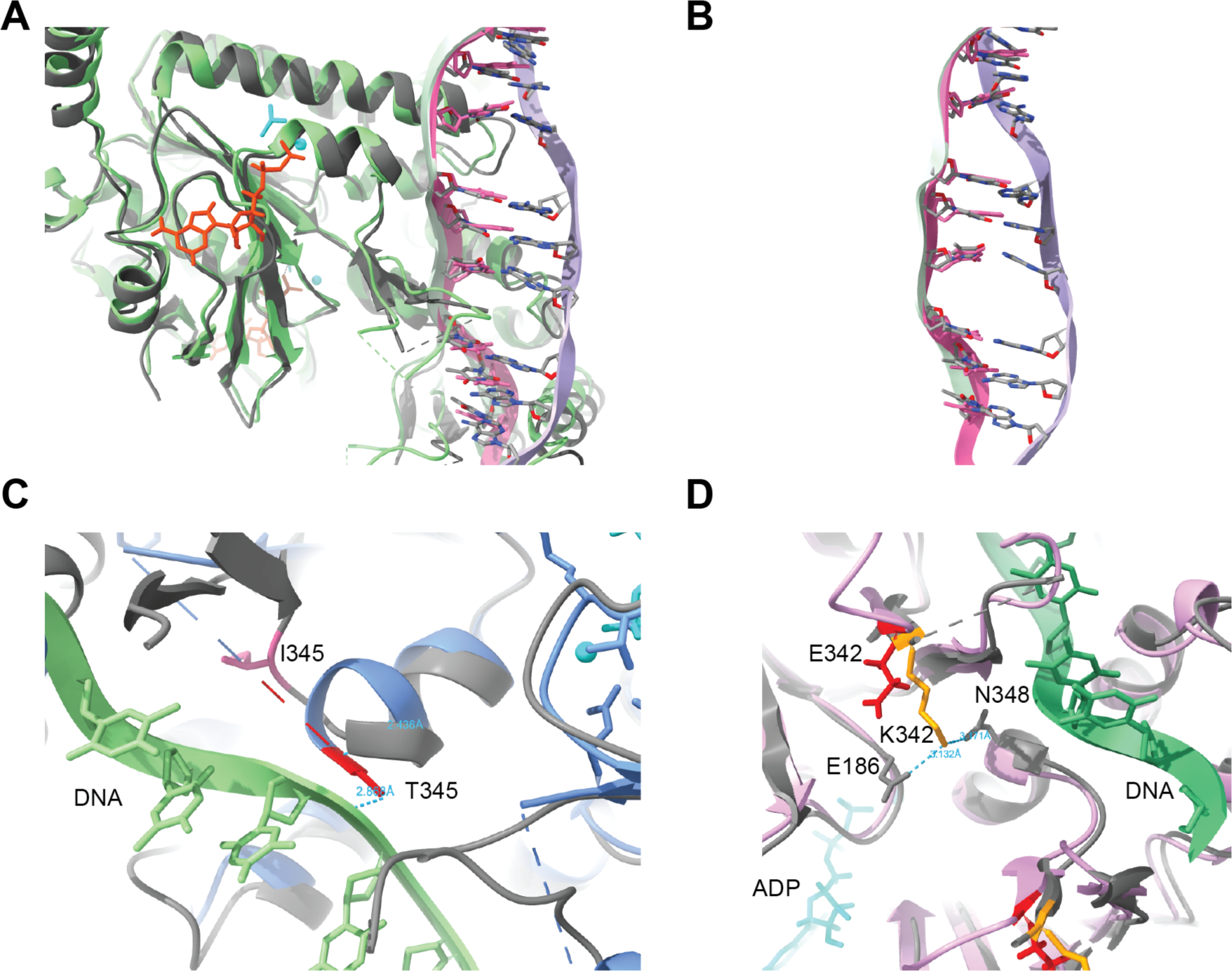
The contacts between Rad51 and DNA, as well as the positions of the DNA backbone and bases, are highly conserved between yeast Rad51-ssDNA and human RAD51-dsDNA complexes. (**A**) Overlay of yeast Rad51 (gray)-ssDNA (pink) model (this study, gray) and human RAD51-dsDNA cryo-EM structure (PDB: 7eje, green). (**B**) Overlay of ssDNA (pink) with dsDNA (green & purple). (**C**) Overlay of WT Rad51-ssDNA cryo-model (this study, gray) and Rad51-I345T crystal structure (PDB: 1SZP, blue). DNA: green. H-bonds formed with mutant Thr345 were shown. (**D**) Overlay of WT Rad51-ssDNA cryo-model (this study, gray) and predicted Rad51-K342E structure (purple). DNA: green.

Next, we compared our reconstructed Rad51 monomer with the two previously published yeast Rad51 mutant structures (I345T and H352Y). The core domain of four chains from three distinct structures aligns well, with only a minor structural divergence observed in the N-terminal helical bundle. This divergence is caused by different hinge orientations between the top and bottom domains (**Suppl. Fig. 4C**). It is most noticeable between the wild type configuration from this study and the H352Y structure (**Suppl. Fig. 4D**), supporting the dynamic movement of the helical top as a cohesive unit and the flexibility of the hinge.

Rad51-I345T is a gain-of-function mutant that exhibits increased DNA-binding activity while retaining normal ATPase activity *in vitro* and biological function *in vivo* (21). Its crystal structure is characterized by a helical filament with a pitch of 130 Å, which is longer than most active Rad51-DNA filaments imaged by EM studies (6,7,18,33,34,37). However, the Rad51-I345T crystal structure lacks the densities of ssDNA and ATP. As a consequence, it does not provide sufficient information to understand how the mutant interacts with ssDNA to increase its binding affinity (18). To bridge this knowledge gap, we have undertaken a comparison of the Rad51-ssDNA structure from this study with the Rad51-I345T structure to explore the differences and gain a deeper insight into the underlying mechanisms.

In wild type Rad51, Ile345 forms two hydrogen bonds with Asn325 and Gln326, located within a small loop away from ssDNA at distances of 2.94 Å and 3.26 Å respectively (**Suppl. Fig. 5A**). However, the mutant I345T establishes a new hydrogen bond with Ile349 at 2.44 Å. This stabilizes and extends the neighboring α-helix to include Gly346 (**Fig. 6C**). Consequently, I345T gets closer to ssDNA and forms a new hydrogen bond with a modeled DNA base at a distance of 2.87 Å (**Fig. 6C**), providing a structural basis for the gain-of-function in DNA binding. Additionally, the α-helix (α6) comprised of residues Ala147 to Glu156 at the Rad51-Rad51 interface shows an average deviation of about 4.91 Å between wild type and I345T. The deviation amplifies through the hinge to the N-terminal top, resulting in a super extended helical I345T filament with a 130 Å pitch (**Suppl. Fig. 5B**).

### A 3D model facilitating artificial intelligence (AI)-guided mutant structure prediction and analysis

Protein structure prediction systems such as AlphaFold play a vital role in creating molecular models of both wild-type and mutant proteins. The precise location of DNA and nucleotide cofactor in our Rad51 structure, in combination with local amino acid dynamics, enables us to improve the analysis of predicted mutant structures. We conducted an analysis of the AI-predicted structure of the yeast Rad51-K342E mutant. This mutant exhibits significant defects in dsDNA-binding, while demonstrating DNA-independent ATPase activity that is sensitive to ADP inhibition. Furthermore, the K342E mutant forms protein filaments in the absence of DNA (38). Lys342 is distributed along the DNA within the helical filament and is situated between ADP-AlF_3_ and DNA (**Suppl. Fig. 6C, D**). In the wild type filament, Lys342 forms hydrogen bonds with Glu186 at 3.13 Å and Asn348 at 3.17 Å, both of which are lost by the Glu342 mutation (**Fig. 6D**). Asn348 interacts with DNA backbone and Glu186 is within the nucleotide-binding Walker A motif, providing a structural basis for the loss of coordination of nucleotide- and DNA-binding in the K342E mutant, as well as the observed defects in DNA-binding.

### Structural analysis of tumor-derived human RAD51-Q268P and RAD51-Q272L mutant proteins

Some human tumors have been found to exhibit altered expression levels and deleterious mutations of RAD51 (23,39–44). It is of interest to investigate whether the local motion or rigidity of individual amino acids of Rad51 we detected can provide additional insights into cancer-relevant mutants beyond their predicted structures. We used two human lung and kidney tumor-derived RAD51 variants, Q268P and Q272L, as test cases of whether examining the local dynamics of RAD51 would provide a more comprehensive understanding of the mechanisms underlying cancer-associated mutations. First, we used AlphaFold to predict stuctures of RAD51-Q268P and RAD51-Q272L, and compared them to the published wild type RAD51 structure (PDB: 7eje), as shown in **Figure 7A**. Both Gln268 and Gln272 are located in loop 2 between AMPPNP and DNA (**Fig. 7A**). No significant change is detected for secondary and overall structures. The high sequence conservation between yeast and human Rad51 (**Fig. 2D**) allows us to explore the mutational consequences in changing the local conformational rigidity of Gln268 (yeast Gln326) and Gln272 (yeast Gln330), as shown in **Figure 7B**. In the wild type yeast Rad51 filament, Gln326 is fixed by the H-bonds with the DNA-interacting Arg287 and the nucleotide-interacting Glu221, which are abolished by the Gln to Pro mutation (**Fig. 7C**). In contrast, Gln268 in all four human RAD51 structures adopts different conformations and lacks the H-bonds we identified in yeast Rad51 structures, justifying the usage of yeast Rad51 structure in interpreting human RAD51 mutants. Additionally, we predicted the structure of yeast Rad51-Q326P and confirmed its structural consistency with human RAD51-Q268P (**Fig. 7C**). Both mutants lost the coordination between ATP- and DNA-binding, providing a structural basis for the dramatic decrease of human RAD51-Q268P in DNA-binding, DNA-stimulated ATP hydrolysis, and the ability to catalyze plasmid-based DNA strand exchange (23). Similarly, we compared predicted structures of yeast Rad51-Q330L and human RAD51-Q272L with wild type yeast Rad51 structure from this study (**Fig. 7D**). The rigidity of yeast Gln330 (human Gln272) is preserved by its H-bond with Pro341 and ensures the rigidity of yeast Val331 (human Val273) to bind and insert into DNA nucleotides (**Fig. 7D, Suppl. Fig. 6E**). Consequently, the mutation of yeast Gln330 to Leu (human Q272L) disrupts an H-bond that maintains structural rigidity, resulting in bad contacts of yeast Val331 (human Val273) with DNA (**Fig. 7E**). The direct impact on DNA explains why RAD51-Q272L displays more severe defects in ssDNA- and dsDNA-binding compared to RAD51-Q268P. The residual DNA-binding activity supports the significant reduction in plasmid-based DNA strand exchange (23).

**Figure 7.**
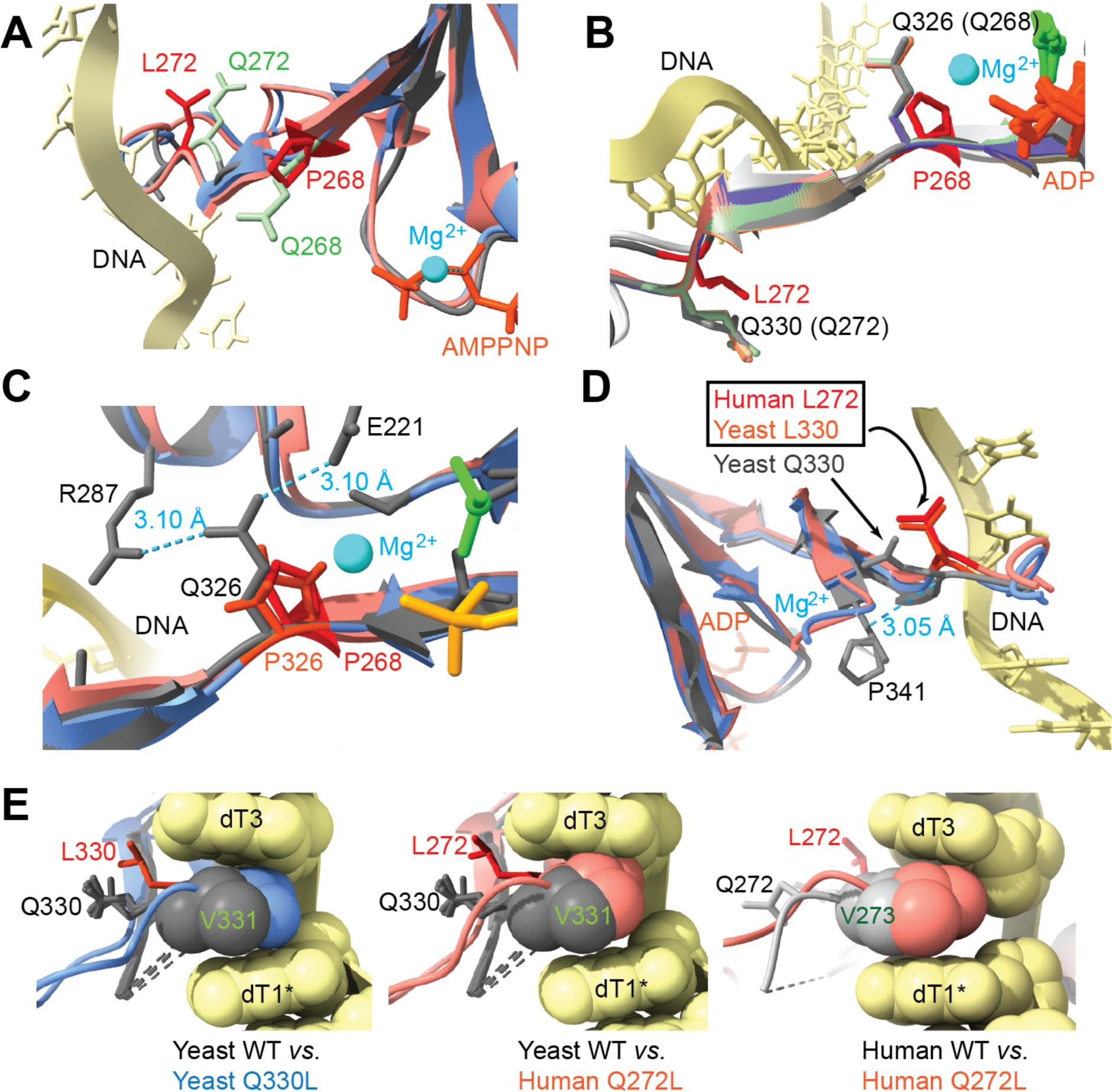
Structural analysis of tumor-derived human RAD51-Q268P and RAD51-Q272L mutants. (**A**) Overlay of predicted RAD51-Q268P (salmon) and RAD51-Q272L (blue) with wild type RAD51 (7ejc, gray) structures. Mutated Pro268 and Leu272 were shown in red and AMPPNP in orange. (**B**) Overlay of predicted RAD51-Q268P (light gray) and RAD51-Q272L (dark gray) with 6 protomers of yeast Rad51 in rainbow colors (this study). Mutated Pro268 and Leu272 were shown in red, while corresponding wild type yeast Gln326 and Gln330 were shown in rainbow colors. ADP: orange; AlF_3_: green. (**C**) Overlay of predicted RAD51-Q268P (salmon) and predicted yeast Rad51-Q326P (blue) with wild type yeast Rad51 (gray). H-bonds of Gln326 with Glu221 and Arg287 in wild type yeast Rad51 structure were shown. Mutated Pro268 in human RAD51 were shown in red and Pro326 in yeast Rad51 were shown in orange. ADP: yellow; AlF_3_: green. (**D**) Overlay of predicted RAD51-Q272L (salmon) and predicted yeast Rad51-Q330L (blue) with wild type yeast Rad51 (gray). H-bond of Gln330 with Pro341 in wild type yeast Rad51 structure was shown. Mutated Leu272 in human RAD51 were shown in red and Leu330 in yeast Rad51 were shown in orange. In **A**-**D**, DNA: yellow; Mg^2+^: cyan. (**E**) Illustration of defective interaction of Val273 in the predicted structure of RAD51-Q272L and Val331 in yeast Rad51-Q330L with DNA. DNA in wild type yeast Rad51 structure (this study) was shown as a reference in yellow.

## DISCUSSION

Helical symmetry has been widely imposed on 3D reconstructions of the negative-stained or cryo-EM density maps of Rad51 filaments on ssDNA and dsDNA in order to generate different views of a single protomer and thus improve structural resolution (7,33,45). For the first time, we performed single-particle analysis without imposed helical symmetry and reconstructed a high-resolution map of Rad51-ssDNA filaments at 2.35 Å. The molecular model of the filaments, captured with the bound ligand ADP-AlF_3_, is consistent with previous cryo-EM studies of human RAD51 filaments with DNA in a high-affinity state (33,34). Comparisons of individually reconstructed protomers lead to the discovery of local structural movements of amino acid side chains within the overall highly conserved and rigid Rad51 nucleoprotein filaments. The observation that these residues in each protomer could be iteratively refined to obtain high-quality density maps indicates the high probabilities of these residues holding the corresponding conformation in the short repetitive filament segment with a directionality. We propose that the initial binding of Rad51 onto the ssDNA backbone is mediated through the positively charged channel (**Suppl. Fig. 3A, B**). The binding process of RAD51 occurs in a DNA sequence-independent manner and is a rapid process that leads to conformational changes in the Loop1 and Loop2 regions. Unlike the flexible Arg293, Leu296 and Val331 are rigid enough to insert between DNA bases and organize the bases into groups of triplet bases. When comparing the structures of human Rad51-ssDNA and RAD51-dsDNA, Arg235 and Asp274 exhibit a swing towards the complementary DNA strand (**Suppl. Fig. 3E, F**). This suggests that the flexible Loop1 and Loop2 play a crucial role in engaging homologous DNA for base pairing and exchange. In brief, the tight binding of the DNA backbone is responsible for the sequence non-specific binding of Rad51. Meanwhile, the subsequent dynamic movements of the amino acids in Loop1 and Loop2 dictate the triplet formation required for homology search and pairing. These local yet dynamic structural changes are exhibited by the protomers in the Rad51-ssDNA filament, akin to a timelapse in frames.

Rapid freezing in Cryo-EM preserves target proteins in their native state with an aqueous buffer, providing a unique opportunity to capture enzymes in action. Transient states of enzymes or reaction intermediates could be pinpointed on the condition that the number of single particles are large enough, similar to single molecule studies. Lys191 in the Walker A motif has been widely characterized as the critical catalytic residue for ATP-binding and hydrolysis (9,13). Our analysis shows that Lys191 is rigid, and this lack of movement probably reflects the non-hydrolysable feature of ADP-AlF_3_. Lys191 is immobilized by a Mg^2+^ ion and the β-phosphate of the ADP-AlF_3_ (**Fig. 4A**). In addition, Ser192 was identified here to play a facilitating role in ATP hydrolysis by pulling AlF_3_ away from ADP (**Fig. 4B**). This action is presumed to work with Lys191 to break the high-energy phosphoanhydride bonds in ATP. Similarly, we noticed the same pattern for human RAD51 that Thr134 displays a pulling action with three different conformations while Lys133 almost remains the same conformation (**Suppl. Fig. 4C, D**). The role of Ser192 in ATP hydrolysis is supported by the consistency in yeast and human RAD51, validating local amino acid dynamics we observed.

The comparison of individual Rad51 protomers in the filament also reveals local amino acid movements at the protein-interaction interface. Phenylalanine (Phe144 in budding yeast Rad51) has been an essential residue in the FxxA motif, discovered as the protein-interaction motif in the BRC repeats of BRCA2 and yeast/human Rad51 (46). As shown in **Figure 5B**, Phe144 sits at the protomer-protomer interface with Val145, Thr146, and Ala147, and all four residues remain tightly bound with not even the slightest of movements of side chains. Interestingly, phenylalanine at other positions also tends to be rigid with no local movements, whether in α-helices or flexible loops. The four amino acids in the false-positive FxxA site, 317-FGVA-320, are located in a similar flexible loop but display no side chain movements. They are buried inside the Rad51 protomer and unlikely involved in self- or hetero-protein interactions of Rad51 (data not shown). The other example is the acidic patch of Rad51 involved in protein interactions with Rad52, Rad55-Rad57, and Rad54 (36). The three responsible aspartates, D239, D241, and D242, are dynamic, accommodating a sensing configuration to engage other protein factors (**Fig. 5D**). Similarly, the amino acids in the vicinity but interacting with neighboring Rad51 protomer show minimal movement without alternative conformations (**Fig. 5E**). Thus, analysis of amino acid dynamics might be effective to identify the key residues in the target motifs for self- and hetero-interactions.

The near atomic-resolution map in this study supports a high-quality fit of a long stretch of ssDNA and the nucleotide cofactor, ADP-AlF_3_ (**Fig. 2**), adding new levels of structural details of the Rad51 nucleoprotein filament. The structural comparisons of the wild type Rad51 with the previously published structures of the I345T and H352Y mutants (18,20) reveal the altered interactions of DNA with a single amino acid change and uncover the mechanism behind their gain-of-function mutant behaviors (**Fig. 6C**). The ATP-binding pocket alignment of all three yeast Rad51 protein species reveals a noteworthy general movement of critical residues towards the pocket, as illustrated in **Figure 8A**. The I345T structure did not detect ATPγS and ssDNA while the H352Y structure does not contain nucleotide and ssDNA. The inward orientation of F187, R188, S192, and Q326 in I345T and H352Y in “no nucleotide” conformation may reflect the transitional state of Rad51 between ADP-release and new ATP-binding (**Fig. 8B**). This configuration might partly constitute the structural deviation between the wild-type Rad51 and I345T/H352Y, particularly in helical pitch (**Suppl. Fig. 5B**).

**Figure 8.**
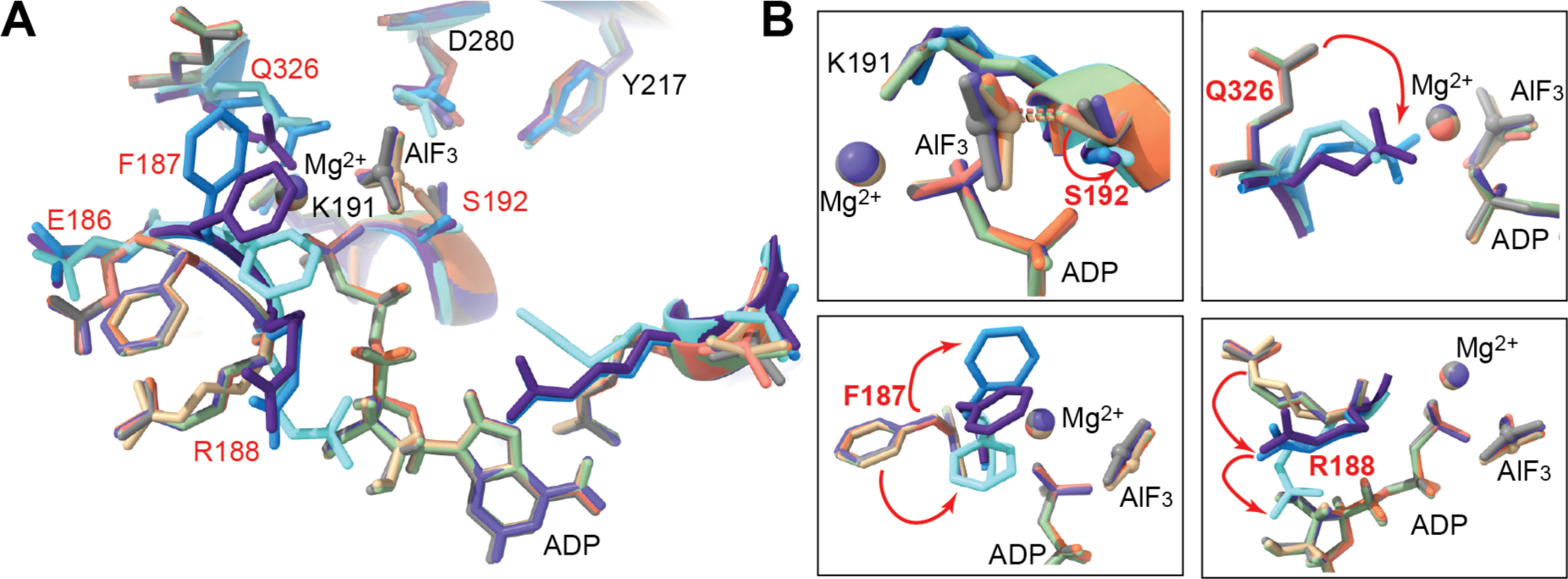
Different configurations of the ATP-binding pocket of the Rad51 protein in its wild-type state and two mutant structures: Rad51-I345T and Rad51-H352Y. (**A**) The ATP-binding pockets from six wild-type Rad51 protomers (this study, rainbow) are overlaid with Chain A (light blue) and Chain D (purple) from Rad51-I345T crystal structure (PDB: 1SZP), and Rad51-H352Y crystal structure (PDB: 3LDA, cyan). (**B**) The orientation of Phe187, Arg188, Ser192, and Gln326 in Rad51-I345T (light blue and purple) and Rad51-H352Y (cyan) are compared to wild-type Rad51 (rainbow).

For the first time, we detected the local dynamics of amino acids of yeast Rad51 involved in binding ATP, DNA, and proteins. We used this information to analyze the structural changes of yeast Rad51-I345T and Rad51-K342E, as well as human RAD51-Q268P and RAD51-Q272L, and provided satisfying explanations of their biochemical defects. We also created a structure of Rad51-dsDNA with modeled-in dsDNA. We envision that the Rad51-dsDNA filament would be useful to analyze certain Rad51 mutant filaments formed on dsDNA, considering the highly conserved positions of DNA in Rad51-ssDNA and RAD51-dsDNA filaments (**Fig. 6A, B**). In summary, our study has provided the first high-resolution structure of the yeast Rad51 filament with detailed information on Rad51-ATP and Rad51-DNA interaction. Additionally, we captured the local dynamics of Rad51 amino acids, which are expected to contribute to the understanding of RAD51 filament formation in HR repair.

## Data Availability

The density maps and data have been deposited in the Electron Microscopy Data Bank under accession codes EMD-44104 and EMD-44105. The atomic coordinates and data have been deposited in the Protein Data Bank under accession code 9B2D.

## Acknowledgments

We thank Phoebe Rice and Scott Morrical for their helpful discussion and comments on the manuscript. The yeast Rad51 expression system was a kind gift from Patrick Sung. We thank Dr. Fei Guo for his technical assistance in grid preparation and data collection. We thank Dr. Wah Chiu (Stanford University) for his technical support and help. Large cryo-EM datasets were collected at the Stanford-SLAC Cryo-EM Center (S^2^C^2^). The content is solely the responsibility of the authors and does not necessarily represent the official views of the National Institutes of Health. The authors would also like to thank the following S^2^C^2^ personnel for their invaluable support and assistance in data collection: Chensong Zhang and Ian Fries.

## Author Contributions

JL and SG conducted cryo-EM sample preparation and data collection; JL performed cryo-EM image analysis and model building; JL, SG, and WDH designed experiments and interpreted results; JL and WDH designed the study; JL, SG, and WDH wrote the manuscript.

